# Effect of cellular de-differentiation on the dynamics and evolution of tissue and tumor cells in mathematical models with feedback regulation

**DOI:** 10.1101/204214

**Authors:** Dominik Wodarz

## Abstract

Tissues are maintained by adult stem cells that self-renew and also differentiate into functioning tissue cells. Homeostasis is achieved by a set of complex mechanisms that involve regulatory feedback loops. Similarly, tumors are believed to be maintained by a minority population of cancer stem cells, while the bulk of the tumor is made up of more differentiated cells, and there is indication that some of the feedback loops that operate in tissues continue to be functional in tumors. Mathematical models of such tissue hierarchies, including feedback loops, have been analyzed in a variety of different contexts. Apart from stem cells giving rise to differentiated cells, it has also been observed that more differentiated cells can de-differentiate into stem cells, both in healthy tissue and tumors, aspects of which have also been investigated mathematically. This paper analyses the effect of de-differentiation on the basic and evolutionary dynamics of cells in the context of tissue hierarchy models that include negative feedback regulation of the cell populations. The models predict that in the presence of dedifferentiation, the fixation probability of a neutral mutant is lower than in its absence. Therefore, if de-differentiation occurs, a mutant with identical parameters compared to the wild-type cell population behaves like a disadvantageous mutant. Similarly, the process of de-differentiation is found to lower the fixation probability of an advantageous mutant. These results indicate that the presence of de-differentiation can lower the rates of tumor initiation and progression in the context of the models considered here.

## 1. Introduction

Healthy tissue is characterized by a specific hierarchical structure, where adult stem cells are responsible for tissue maintenance and give rise to a population of terminally differentiated cells that perform their designated function (Weissman, 2000). Typically, adult stem cells have self-renewal capacity, but also differentiate into transit amplifying cells, which in turn have limited self-renewal capacity and give rise to terminally differentiated cells. Data suggest that tissue homeostasis is maintained through the existence of specific positive and negative feedback loops that influence the stem cell numbers and tissue size (Watt and Hogan, 2000).

Tumors arise from healthy tissues and are thought to maintain some of the hierarchy characteristic of healthy tissue (Jordan et al., 2006; Reya et al., 2001; Visvader and Lindeman, 2012). Hence, it is thought that tumors are maintained by so called cancer stem cells (CSC), and that the majority of the tumor is made up of more differentiated cancer cells that have limited ability to maintain the existence of the tumor. Similarly, it has been suggested that feedback loops which regulate healthy tissue homeostasis are maintained to a certain extent in tumors (Rodriguez-Brenes et al., 2011), and observed tumor growth patterns support this notion (Rodriguez-Brenes et al., 2013b). While the existence of feedback control in tumors has not been much investigated experimentally, there is mounting evidence that feedback regulatory mechanisms play a role in tumor growth dynamics (Kurtova et al., 2015; Rodriguez-Brenes et al., 2017). It has been argued that remaining feedback control mechanisms in tumors are responsible for the often described logistic or Gompertzian growth patterns, where cell populations initially grow, but subsequently stabilize around an apparent equilibrium for a certain period of time before progressing further (Rodriguez-Brenes et al., 2011).

Both healthy tissue as well as tumor cell dynamics have been studied with mathematical models in the context of hierarchically structured cell populations (Anderson and Quaranta, 2008; Enderling and Hahnfeldt, 2011; Enderling et al., 2013; Enderling et al., 2007; Glauche et al., 2007; Komarova and van den Driessche, 2017; Lander et al., 2009; Lo et al., 2009; Marciniak-Czochra et al., 2009; Michor, 2008; Rodriguez-Brenes et al., 2013a; Roeder and Loeffler, 2002; Roederer et al., 2006; Stiehl and Marciniak-Czochra, 2011; Sun and Komarova, 2015; Werner et al., 2011; Werner et al., 2016; Yang et al., 2015). These approaches typically assume a uni-directional differentiation pathway, where stem cells give rise to transit amplifying cells, which in turn give rise to differentiated cells. Recent data, however, indicate that a degree of phenotypic plasticity can occur, and that more differentiated cells can de-differentiate into stem cells, both in tumor cells and in healthy tissue (Cabrera et al., 2015; Chaffer et al., 2013; Chaffer et al., 2011; Dorantes-Acosta and Pelayo, 2012; Gupta et al., 2011; Huels and Sansom, 2015; Kreso and Dick, 2014; Li and Laterra, 2012; Mani et al., 2008; Marjanovic et al., 2013; Philpott and Winton, 2014; Scheel and Weinberg, 2011; Tata et al., 2013). In healthy tissue, de-differentiation tends to be observed during the process of tissue regeneration (Desai et al., 2014; Stange et al., 2013; Yanger and Stanger, 2014; Yanger et al., 2013), probably because at homeostasis, the de-differentiation process might be suppressed through negative feedback resulting from the contact of more differentiated cells with stem cells (Tata et al., 2013). A limited number of mathematical modeling studies have taken into account the concept of phenotypic plasticity in the context of cancer (Jilkine and Gutenkunst, 2014; Kaveh et al., 2016; Leder et al., 2010; Roeder and Loeffler, 2002; Shirayeh et al., 2016; Tonekaboni et al., 2017), including an investigation of de-differentiation on the evolution of mutants (Shirayeh et al., 2016). At the same time, however, our understanding about the effect of cell de-differentiation on basic tissue dynamics and cellular evolution remains incomplete. Here, we build on previous mathematical models that take into account tissue hierarchy and regulatory feedback loops to further study the effect of cellular de-differentiation on the dynamics and evolution of cells, both in healthy tissue and in tumors. We start by considering ordinary differential equation models to examine how de-differentiation affects the ability to maintain tissue homeostasis. We then consider stochastic models to investigate the effect of de-differentiation on the evolution of mutant cell populations.

## 2. Basic model of stem-cell driven tissue dynamics

Here, we review a basic model of stem cell-driven tissue dynamics. In the simplest form, this model contains two populations: the stem cells, S, and the more differentiated cells, D. The latter population is assumed to capture both the transit amplifying and differentiated cells. This model can correspond to either healthy tissue cells or to tumor cell populations, depending on the scenario to which the model is applied. The model is given by the following set of ordinary differential equations (see also (Lander et al., 2009) for related models).

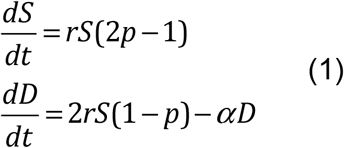

Stem cells are assumed to divide with a rate r. This division results in the generation of 2 daughter stem cells (self-renewal) with a probability p. With a probability 1-p, the division results in the generation of two differentiated cells. Therefore, division is assumed to be symmetric, which leads to either self-renewal or differentiation with given probabilities. Finally, differentiated cells are assumed to die with a rate α. In this model, two outcomes are possible. If the self-renewal probability p<0.5, the cell populations go extinct. In contrast, if the self-renewal probability of cancer stem cells p>0.5, exponential growth is observed. An equilibrium is not possible in this model except in the case when p=0.5.

## 3. Stabilization through feedback

In previous modeling approaches, it has been assumed that differentiated cells secrete negative feedback factors that (i) reduce the rate of cell division, and (ii) reduce the self-renewal probability of stem cells, based on experimental observations (Konstorum et al., 2016; Lander et al., 2009; Lo et al., 2009; Rodriguez-Brenes et al., 2011; Rodriguez-Brenes et al., 2017). In the model, this can be expressed by replacing the division rate and the self-renewal probability with the terms r = r’/(1+h_1_D^k1^) and p = p’/(1+h_2_D^k2^), where r’ and p’ are the basic division rate and the self-renewal probability of stem cells in the absence of any feedback. According to these assumptions, as the overall number of stem cells increases, the number of differentiated cells also rises, resulting in inhibition of stem cell self-renewal. Instead, more stem cell divisions result in differentiation and hence eventually in death of the cells. In this model, persistence of the cell populations requires that p>0.5, in which case the system converges to the following stable equilibrium.

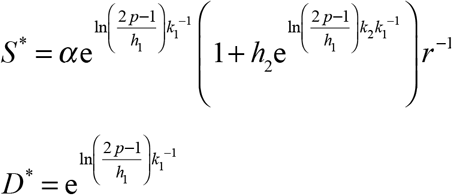

## 4. The simplest model with stem cell plasticity

The above model can be extended to include the process of de-differentiation in the following way:

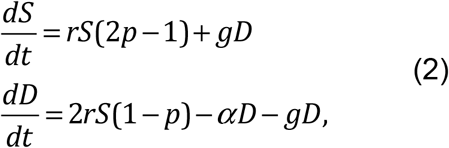

Where r = r’/(1+h_1_D^k1^) and p = p’/(1+h_2_D^k2^). In addition to the basic processes already described, differentiated cells are assumed to de-differentiate into stem cells with a constant rate g. For now, this model assumes that de-differentiation is not subject to any feedback, which will be introduced later.

In this model, persistence of the cell populations is observed if 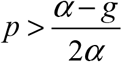. In contrast to the model without plasticity, persistence of cells is now possible for p<0.5. If the rate of de-differentiation is greater than the death rate of differentiated cells (g>α), the condition for the persistence of cells becomes p≥0. That is, the cell populations can grow even in the absence of self-renewal, because the generation of new stem cells from differentiated cells is sufficient to drive a population increase. If the condition for the persistence of the cell populations is fulfilled, and if the rate of dedifferentiation g<α, the system converges to the following stable equilibrium.

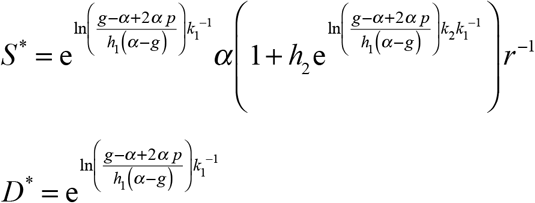

If g>α, on the other hand, no stable internal equilibrium exists, and the cell populations grow unbounded towards infinity. This growth is slower than exponential, driven by the feedback loops that continue to operate in the cell population. Therefore, we note that if the de-differentiation rate is greater than the death rate of differentiated cells, the negative feedback loops on stem cell divisions become unable to prevent unbounded growth of cells, i.e. to maintain a degree of homeostasis. We further note that this is independent of the parameters that describe the strength of the negative feedback on stem cell self-renewal, h and k. That is, if g>α, an equilibrium and homeostasis are impossible, no matter how strong the degree of negative feedback on self-renewal. This corresponds to the parameter regime in which stem cell self-renewal is not strictly required for cell growth, due to the replenishment of stem cells through de-differentiation. This becomes increasingly relevant for lower death rates of differentiated cells, α.

## 5. Feedback on the rate of de-differentiation

The above model assumed that differentiated cells can de-differentiate to become stem cells with a constant rate, i.e. this process did not include feedback regulation. This could be an appropriate assumption for a cancerous state. Data from secretory cells in mice (Tata et al., 2013), however, indicate (i) that dedifferentiation can occur in healthy tissue in vivo, and (ii) that the process of dedifferentiation might be subject to negative feedback control. Contact of a differentiated cell with a stem cell prevented the occurrence of de-differentiation (Tata et al., 2013). This negative feedback can be incorporated into our model by writing g = g’/(1+h_3_S^k3^), where g’ denotes the rate of de-differentiation in the absence of feedback. Negative feedback on the de-differentiation process enables the existence of a stable equilibrium even if the basic de-differentiation rate is relatively high such that g’>α. The equilibrium expressions for this outcome are too complex to write down here. The cell populations persist at equilibrium if 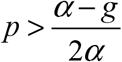.

In the following, we focus on the contribution of self-renewal and dedifferentiation processes to the rate of tissue re-generation and to the protection against uncontrolled growth. We compare two parameter regions: (i) In the first case, either stem cell self-renewal alone, or the process of de-differentiation alone, can maintain the tissue cell population. In terms of model parameters, this translates into p>0.5 and g>α. (ii) The second case assumes the opposite, where both self-renewal and de-differentiation are required to work in concert for tissue maintenance, i.e. p<0.5 and g<α.

If either self-renewal alone or de-differentiation alone can maintain the tissue, the rate of tissue regeneration upon damage is relatively fast (Figure 1A, blue lines). In addition, if one of these mechanisms fails and ceases to contribute to stem cell expansion, the tissue can still be maintained by the second mechanism due to redundancy. At the same time, however, escape from just one of the feedback loops that maintain homeostasis (feedback on self-renewal or feedback on de-differentiation) is sufficient to result in uncontrolled growth (Figure 2A). For example, if the cell acquires a mutation to escape feedback on de-differentiation, uncontrolled growth is observed no matter how strong the feedback on the probability of stem cell self renewal. The reason is that de-differentiation provides a separate pathway for amplifying the number of tumor stem cells, which can thus be achieved even without self-renewal. If tissue maintenance requires a collaboration of both self-renewal and de-differentiation, the rate of tissue regeneration is slower, because the total expansion capacity of the stem cells is reduced (Figure 1A, red lines). Further, if one of these two mechanisms fails, the other is unable to compensate, leading to an inability to maintain the tissue. At the same time, however, simultaneous escape from both the feedback on self-renewal and on de-differentiation is required to achieve uncontrolled growth (Figure 2B). The reason is that expansion of the stem cell population is not possible with either stem cell self-renewal or with dedifferentiation alone. Hence, this suggests the existence of a tradeoff between the rate at which tissue can regenerate in response to damage (and the robustness of tissue maintenance), and the number of steps that are required for cells to grow uncontrolled.

**Figure 1.**
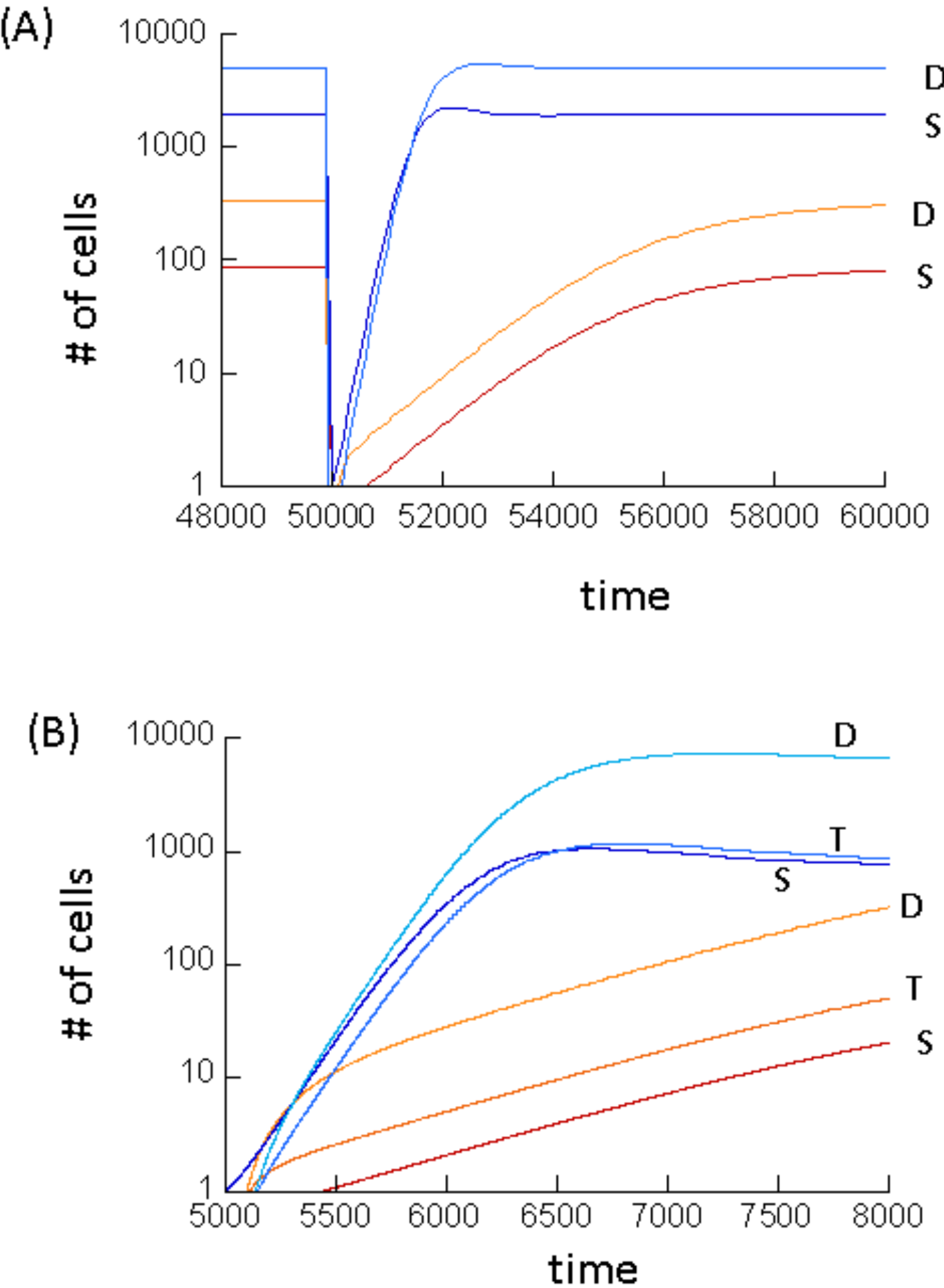
Tissue regeneration dynamics predicted by the mathematical models, following depletion of all cell types. (A) Model (2) with negative feedback on dedifferentiation. The red lines show stem and differentiated cell dynamics arising from simulations assuming that both stem cell self-renewal and de-differentiation need to collaborate to maintain the tissue, i.e. p<0.5 and g’<a. The blue lines assume that either mechanism alone can maintain the tissue, i.e. p>0.5 and g’>a. Tissue depletion was modeled by setting S=1 and D0. Parameters were chosen as follows. r’=0.01; α=0.0025; h_1_=h_2_=0.0001;h_3_=0.01; k_1_=k_2_=k_3_=1. For red lines p’=0.0.35; g=0.0015. For blue lines, lines p’=0.7; g=0.0035. (B) Same, but with model (3), taking into account transit amplifying cells. Tissue depletion was modeled by setting S=1, T=0, D=0. Parameters were chosen as follows. r’1=0.01; r’2=0.02; p’2=0.4; α=0.0025, h1=h2=h3=h4=0.0001; h5=0.01; k_1_=k_2_=k_3_=k_4_=k_5_=1. For red lines p_1_’=0.0.35; q=0.15. For blue lines, lines p_1_’=0.7; q=0.4.

**Figure 2.**
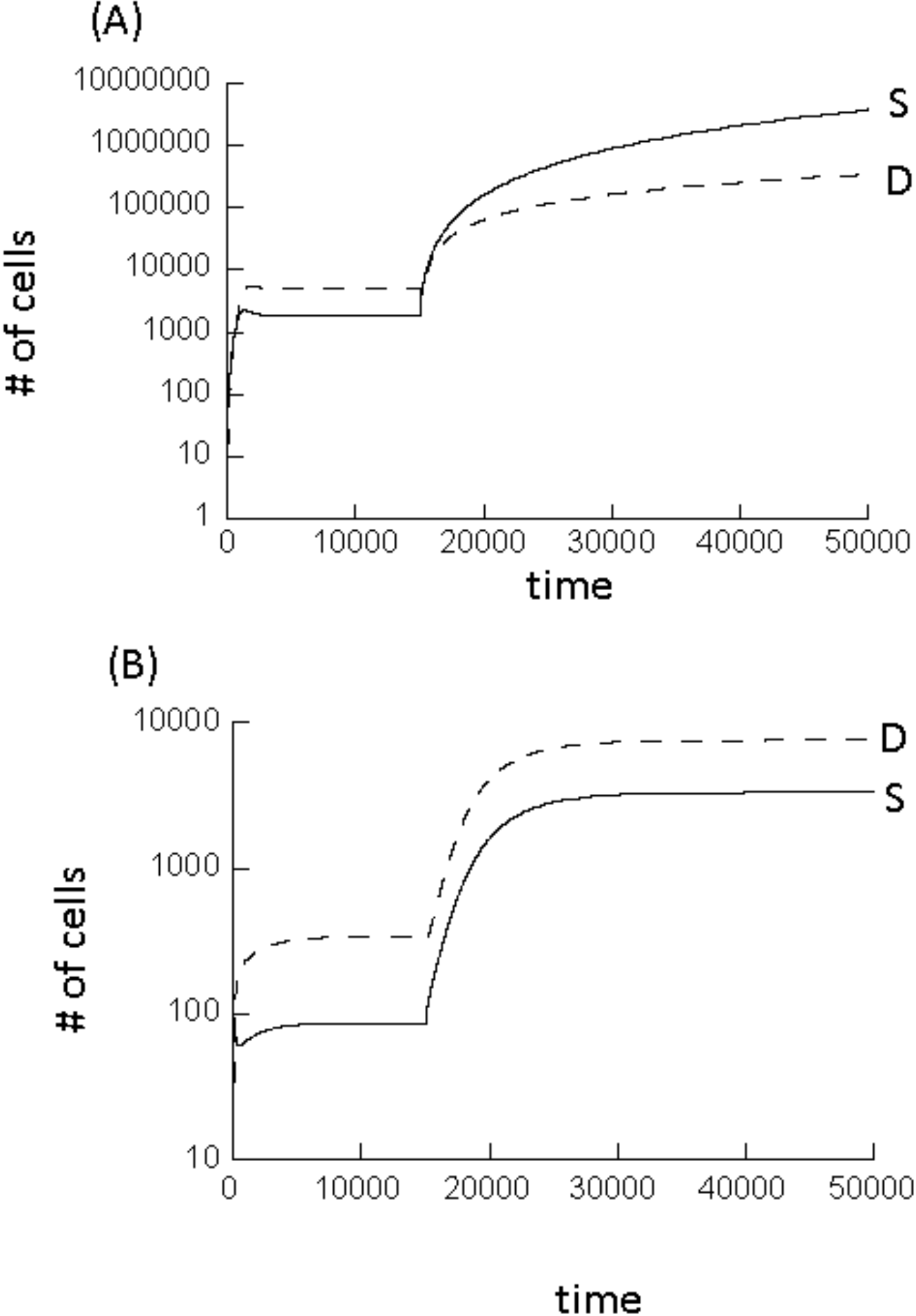
Computer simulation of model (2) with negative feedback on dedifferentiation, assuming that at a specific time point, the feedback on the dedifferentiation process is lost. (A) This simulation assumed that either stem cell self-renewal alone or de-differentiation alone can drive stem cell expansion. Hence, loss of negative feedback on de-differentiation results in uncontrolled growth of the cell populations. (B) This simulation assumes that a combination of self-renewal and de-differentiation is required to drive stem cell expansion. In this case, loss of negative feedback on de-differentiation results in an increase of the equilibrium population sizes, but not in uncontrolled growth. Parameters were chosen as follows. (A) r’=0.01; p’=0.7; α=0.0025; g=0.0035; h_1_=h_2_=0.0001; h_3_=0.01; k_1_=k_2_=k_3_=1. To simulate escape from negative feedback on dedifferentiation, the simulation set h_3_=0. (B) Same but p’=0.35 and g=0.0015.

## 6. Model with transit amplifying cells

Here, we consider a model with more biological complexity, including a population of transit amplifying cells (T) in addition to stem cells (S) and differentiated cells (D). The model is given by the following equations.

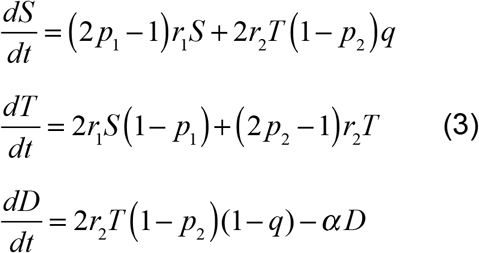

Stem cell division is modeled in the same way as before. Cell division occurs with a rate r_1_, and the division results in self-renewal with a probability p_1_ and in differentiation with a probability 1-p_1_. In the current model, however, differentiation results in the generation of transit amplifying (TA) cells. TA cells can divide with a rate r_2_. This division results in self-renewal with a probability p_2_, and in a differentiation event with a rate 1-p_2_. It is reasonable to assume p_2_<0.5, i.e. TA cells cannot maintain the tissue by themselves. The differentiation of TA cells can lead to two outcomes: with a probability 1-q, a terminally differentiated cell is generated. With a probability q, a de-differentiation event occurs and a stem cell is generated. In contrast to the previous model, the de-differentiation event is now coupled to cell division, since it occurs in the TA compartment. We assume the same types of negative feedback as before. That is, differentiated cells secrete factors that inhibit the rate of stem cell division and the probability of stem cell self-renewal. Thus, as before, we have r_1_ = r_1_’/(1+h_1_D^k1^) and p_1_ = p_1_’/(1+h_2_D^k2^). In the same way, the differentiated cells are assumed to feed back onto the division dynamics of transit amplifying cells, such that r_2_ = r_2_’/(1+h_3_D^k3^) and p_2_ = p_2_’/(1+h_4_D^k4^). In this model, the cell populations persist if 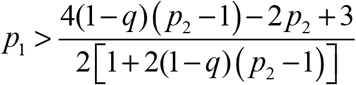. In this case, two types of behaviors are possible. If the probability of de-differentiation during a differentiating TA division q<0.25, the system converges towards a stable equilibrium, which is too complex to write down. If, however, q>0.25, then unbounded growth is observed and negative feedback on stem cell self-renewal fails to contain the number of cells, again irrespective of the strength of the negative feedback on stem cell division (parameters h and k).

As before, we can incorporate negative feedback of stem cells on the de-differentiation probability, writing q = q’/(1+h_5_S^k5^), where q’ denotes the rate of de-differentiation in the absence of feedback. This stabilizes the system even if q>0.25 (again, equilibrium expressions are too complex to be written down). This again presents the opportunity to drive cell expansion via two pathways, i.e. via self-renewal and via de-differentiation. As in the simpler model, if the tissue cannot be maintained by either process alone (p_1_<0.5 and q<0.25), then the cells need to escape both feedback processes simultaneously to achieve uncontrolled growth. At the same time, however, the rate of cell re-population following tissue injury is relatively slow (Figure 1B, red lines), and break-down of only one of those mechanisms results in failure to maintain the tissue. If, in contrast, either stem cell self-renewal or de-differentiation alone can maintain the tissue, then escape from just one feedback mechanism leads to uncontrolled cellular growth, although the rate of cell re-population following tissue injury is faster (Figure 1B, blue lines), and there is redundancy, making tissue maintenance more robust. This again points towards a tradeoff between the rate of tissue repair / robustness of tissue maintenance, and the potential to protect against uncontrolled cellular growth.

## 7. De-differentiation and evolutionary dynamics

Here, we investigate how the process of de-differentiation influences the evolutionary dynamics of mutant cell clones in the context of a resident cell population at equilibrium, maintained by feedback control. For simplicity, we start by considering model (2) with only stem and differentiated cells, and do not take into account feedback on de-differentiation. Two populations are modeled. The “wild-type” populations are denoted by S_1_ and D_1_, and the “mutant” populations are denoted by S_2_ and D_2_, respectively. It is assumed that differentiated cells of both types can secrete feedback factors that regulate the rate of cell division and the probability of self-renewal, and that the stem cells of both types are susceptible to those feedback factors, irrespective of the cell of origin. This introduces competition among the two cell strains mediated through negative feedback. To study the evolutionary dynamics of mutants, stochastic models have to be used, and we peformed Gillespie simulations (Gillespie, 1976) of model (2).

It is assumed that the mutant is neutral, i.e. that it is characterized by the same parameter values as the wild-type. The wild-type population will be assumed to persist at equilibrium. Into this population, a single mutant stem cell, S_2_, is placed. We numerically determined the fixation probability of the mutant cell population. To do so, the simulation was run repeatedly, recording the number of realizations during which the mutant fixated, and those during which the mutant cell clone went extinct. We start by setting g=0, i.e. no de-differentiation occurs in the cell population. In this case, the fixation probability is given by 1/S_1_^*^, where S_1_^*^ denotes the equilibrium number of wild-type stem cells (Figure 3A). This is in accord with basic evolutionary theory (Ewens, 2004; Hartl and Clark, 1997; Nei, 1975), where the fixation probability of one neutral mutant is given by the inverse of the total populations size. Because only stem cells contribute to cellular reproduction, the fixation probability is given by the inverse of the total stem cell population. Next, we assumed g>0, that is, de-differentiation occurs with a rate, g. Now, the fixation probability of the mutant becomes lower than 1/S_1_^*^ (Figure 3A). The larger the value of g, the lower the fixation probability becomes relative to the value of 1/S_1_^*^ (Figure 3A). In fact, the fixation probability can become even lower than 1/(S_1_^*^ +D_1_^*^), i.e. lower than the inverse of the total population size (Figure 3A). In other words, even if de-differentiation implies that some of the differentiated cells also contribute to cellular reproduction, the fixation probability is lower than expected for a neutral mutant. Instead, the observed fixation probability suggests that the mutant is in fact disadvantageous, despite identical parameters.

**Figure 3.**
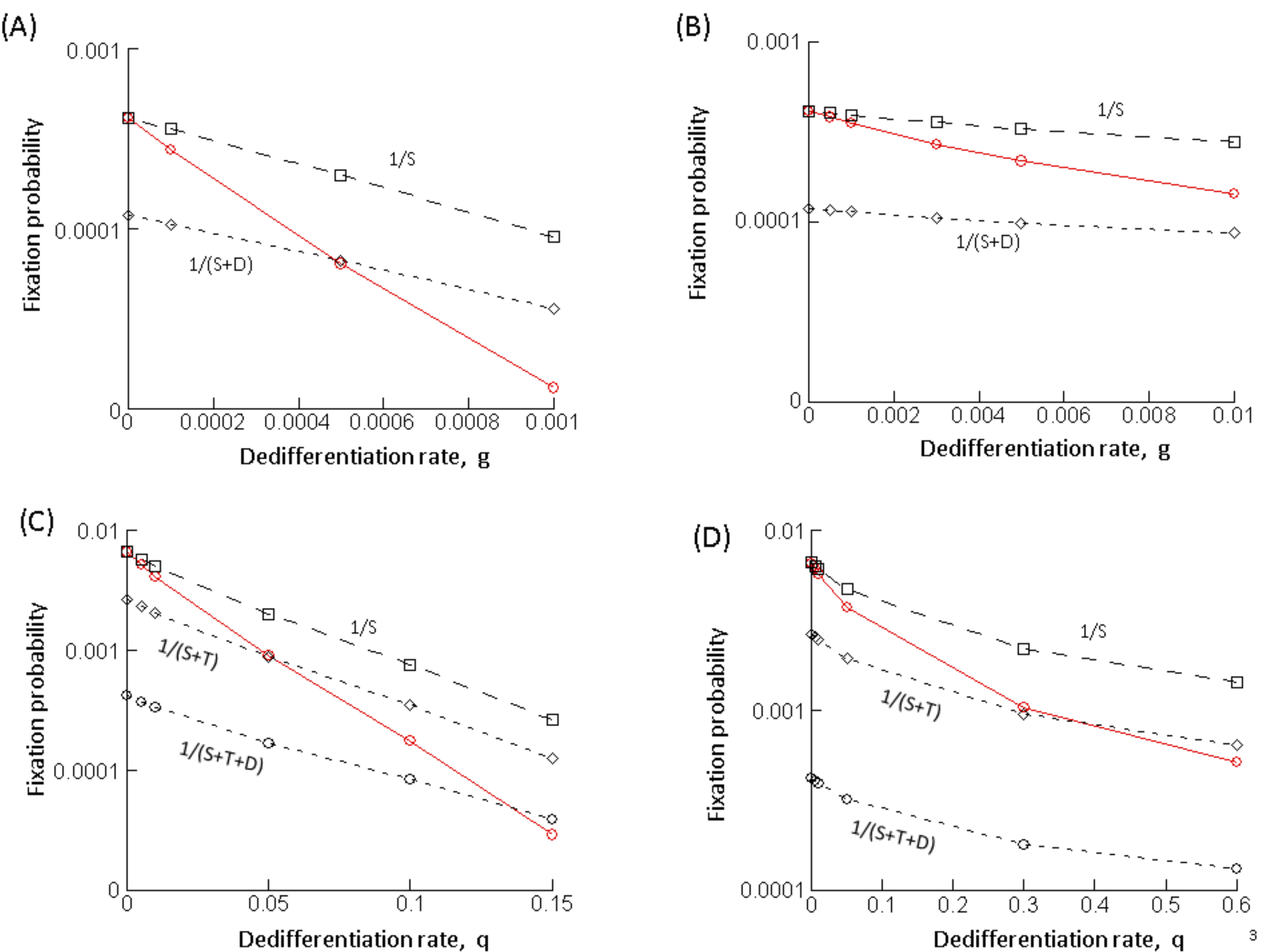
Effect of de-differentiation on the fixation probability of a neutral mutant, determined by computer simulations. Gillespie simulations of the model were run. A single mutant was placed into a resident cell population at equilibrium, and the fraction of realizations that resulted in the fixation of the mutant was recorded. This was done for a case without de-differentiation (g=0 or q=0) and for cases with varying rates of de-differentiation (g or q). This is depicted by the red line in the graphs. The black dashed lines indicate reference values, such as 1/S, the inverse of the equilibrium number of resident stem cells at equilibrium, which according to evolutionary theory, should equal the fixation probability of a neutral mutant. (A) model (2) without feedback on dedifferentiation; r’=0.01; p’=0.8; α=0.0025; h_1_=h_2_=0.0001, k_1_=k_2_=1. (B) model (2) with feedback on de-differentiation; same parameters, and h_3_=0.01, k_3_=1. (C) model (3) without feedback on de-differentiation; r’_1_=0.02; p’_1_=0.6; r’_2_=0.02; p’_2_=0.4, h_1_=h_2_=h_3_=h_4_=0.0001; k_1_=k_2_=k_3_=k_4_=1. (D) model (3) with feedback on dedifferentiation; same parameters, and h_5_=0.01, k_5_=1. Because the mutant was assumed to be neutral, parameters were identical for the resident and mutant cell populations. For each parameter combination, >10^8^ realizations of the simulation were run.

This result can be understood intuitively in the following way. At equilibrium, the total increase of the wild-type stem cell population is governed by two processes: the self-renewal of the stem cells, and the influx from the differentiated cell compartment through the process of de-differentiation. When a single mutant stem cell is introduced into this setting, however, only one of these processes initially contributes to their growth: the self-renewal of the mutant stem cells. Since no differentiated cells initially exist, the de-differentiation process does not contribute to stem cell growth. Hence, at this stage of the dynamics, the total growth rate of the mutant stem cell population is initially lower than that of the wild-type stem cells. Consequently, the mutant experiences an initial disadvantage. As the mutant stem cells build up their differentiated cell compartment, this disadvantage vanishes and the mutant attains neutral properties. This can be observed in simulations of the ODE model (2), see Figure 4. For initial conditions where the wild-type exists at equilibrium, and mutant stem cells are introduced, at first, the number of stem cells declines until the differentiated cell population is established, at which stage the mutant cell population ceases to change in abundance. Overall, this initial and temporary disadvantage of the mutant cell population translates into a fixation probability that is lower than expected for a neutral mutant. These dynamics are not specific to a particular parameter set, but are relevant to all parameter regions in which the differentiated cell population size is abundant relative to the number of stem cells, and in which de-differentiation contributes to sufficiently to stem cell dynamics.

**Figure 4.**
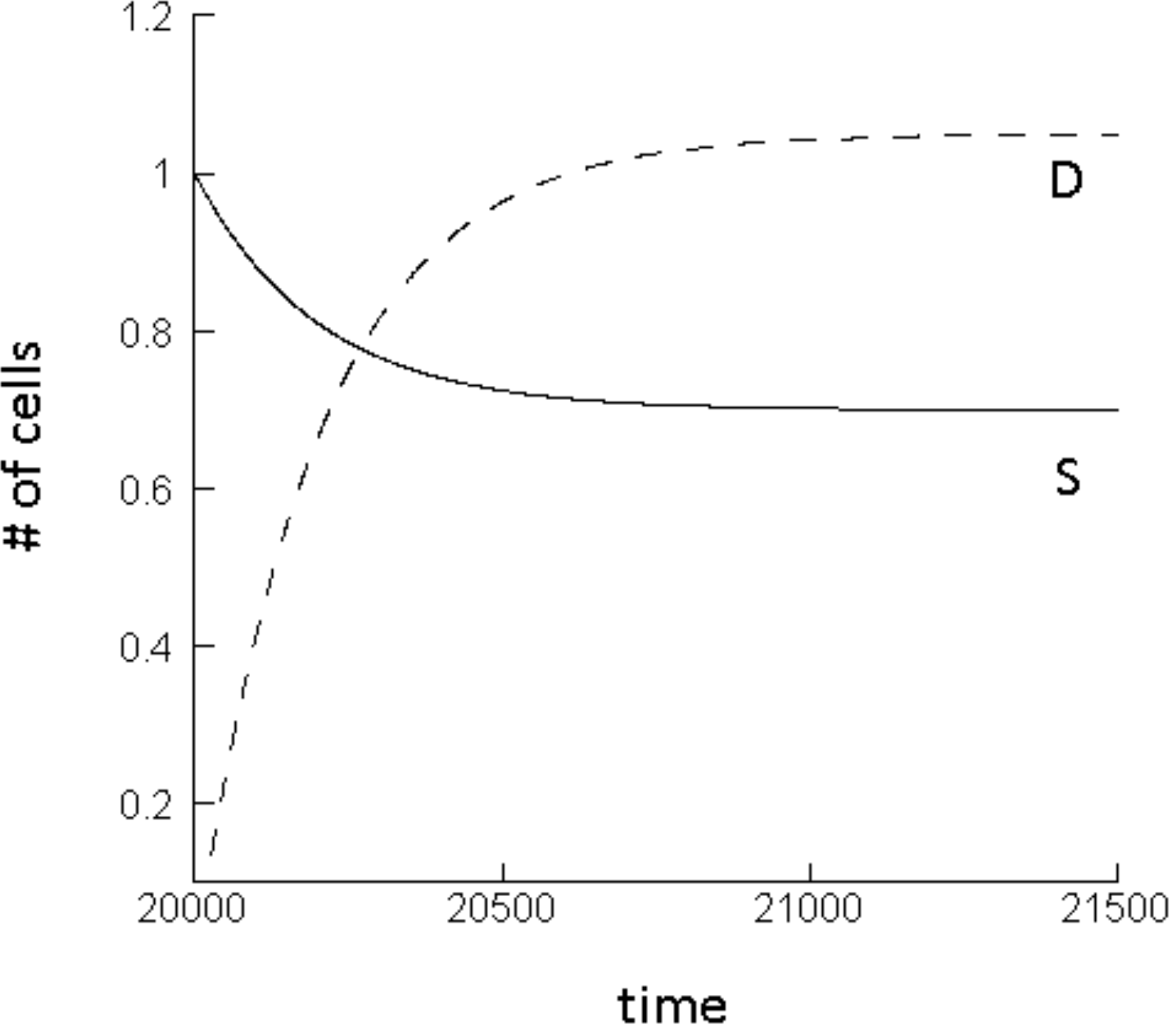
Mutant dynamics predicted by the ODE model (2) without feedback on de-differentiation. The number of mutant stem cells initially declines until the differentiated cell population has built up, at which point the populations converge to a neutrally stable equilibrium. Parameters were chosen as follows. r’=0.01; p’=0.8; α=0.0025; g=0.001; h_1_=h_2_=0.0001; k_1_=k_2_=1.

Similar results are obtained if negative feedback on de-differentiation is assumed to occur, although the effect is smaller (Figure 3B) because the rate of de-differentiation is suppressed by the negative feedback. In addition, results remain robust in the context of model (3) that explicitly takes into account a population of transit amplifying cells (Figure 3 C&D).

The analysis has concentrated on the fixation probability of a neutral mutant. The same kind of result, however, is observed for the fixation probability of an advantageous mutant, which is shown for model (2) without feedback on de-differentiation (Figure 5). It was assumed that the mutant had a larger self-renewal probability, p’, than the wild type, which results in a selective advantage of the mutant in this model (for an analysis of the selective properties of different types of mutants in this kind of model, see (Rodriguez-Brenes et al., 2011)). Again, the process of de-differentiation reduces the fixation probability of the mutant.

**Figure 5.**
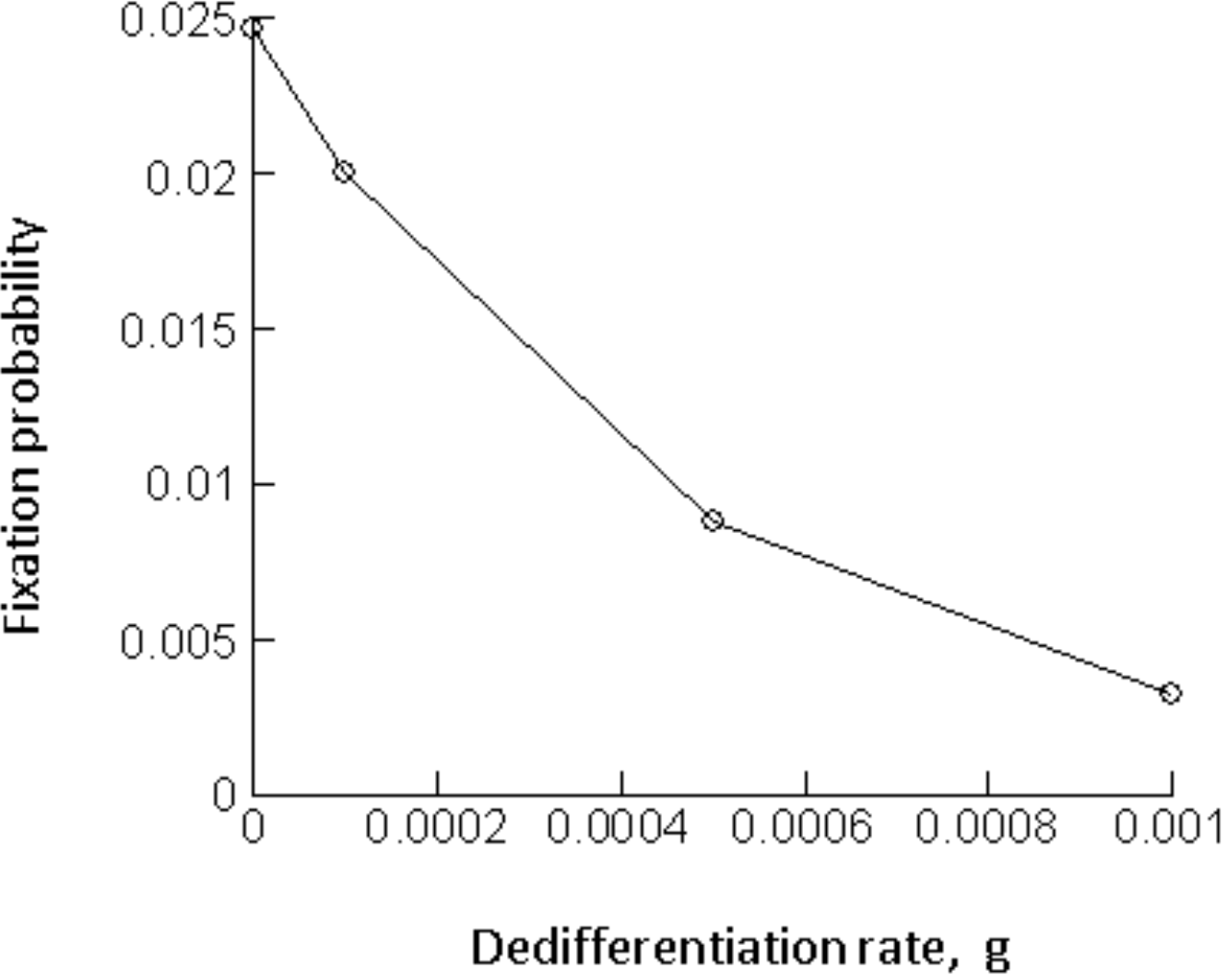
Effect of de-differentiation on the fixation probability of an advantageous mutant, determined by simulations of model (2) without feedback on de-differentiation. Details are the same as in Figure 3. Parameters were chosen as follows. For the resident cellpopulation, r’=0.01; p’=0.8; α=0.0025; h_1_=h_2_=0.0001, k_1_=k_2_=1. The advantageous mutant was characterized by the same parameters, except p’=0.81. For each parameter combination, >10^8^ realizations of the simulation were run.

## 8. Discussion and Conclusion

We used mathematical models to investigate the effect of de-differentiation on basic tissue dynamics as well as on the evolutionary dynamics of cells. Evidence is mounting that de-differentiation occurs in tumor cell populations, and also in healthy tissue, as detailed in the Introduction section. In tumors, it has been suggested that de-differentiation is a process that can fuel tumor growth, progression, and treatment resistance, because it increases the growth rate of the tumor stem cell population, thus rendering growth more robust and increasing resilience against therapeutic interventions (Leder et al., 2010). Kaveh et al (Kaveh et al., 2016) showed that the generation of tumor cells with increased dedifferentiation potential can contribute to their selection, thus enhancing the process of carcinogenesis. Interestingly, a recent paper by Shirayeh et al (Shirayeh et al., 2016) also studied the fixation probability of mutants assuming that both wild-type and mutant cells can de-differentiate, in the context of a Moran process model. They found that the presence of de-differentiation can increase the fixation probability of neutral mutants.

The models presented here, however, argue that the process of dedifferentiation has the opposite effect on the fixation probability of neutral mutants. That is, in the presence of de-differentiation, the fixation probability of a neutral mutant is lower than predicted by neutral evolutionary theory. In other words, in the presence of de-differentiation, a mutant with identical parameters compared to the wild-type acts like a disadvantageous mutant. Similarly, the models suggest that de-differentiation also reduces the fixation probability of advantageous mutants. These results indicate that the presence of dedifferentiation can slow down the emergence of mutants involved in the process of carcinogenesis, and can thus reduce the chance of disease development. This can apply to the development of mutant clones in healthy tissue, but also applies to the progression of already initiated tumors, which also relies on the emergence of mutant clones that overcome relevant selective pressures. In fact, this effect might be more pronounced in tumors than in healthy tissues. According to the model, the reduced fixation probability of mutants is more pronounced if there is no feedback on the process of de-differentiation, which is likely to be the case in tumors but unlikely in healthy tissue. The reason for the reduced fixation probability in the presence of de-differentiation is that mutant clones consist initially mostly of stem cells, and thus do not benefit from stem cell expansion due to de-differentiation. This is in contrast to the established wild-type cell population, in which the number of stem and differentiated cells has equilibrated, and where de-differentiation plays an important role in the dynamics of stem cells.

The reason for the discrepancy between our results and those obtained by Shirayeh et al (Shirayeh et al., 2016) are currently not clear. We considered Gillespie simulations of ODEs that take into account not only tissue hierarchy but also negative feedback processes that regulate the rate and pattern of divisions and differentiations. Stem cell divisions were assumed to be symmetric, giving rise to two progeny stem cells with a probability p, and to two progeny differentiated cells with a probability 1-p. This mechanism is supported by experimental data from human tissues (Nicolas et al., 2007; Simons and Clevers, 2011; Snippert et al., 2010), and this model structure has an established history in the literature (Konstorum et al., 2016; Lander et al., 2009; Lo et al., 2009; Rodriguez-Brenes et al., 2011; Rodriguez-Brenes et al., 2013a; Rodriguez-Brenes et al., 2013b; Rodriguez-Brenes et al., 2015; Rodriguez-Brenes et al., 2017). We considered models with different complexity, taking into account just stem and differentiated cells in the simplest version, but also explicitly taking into account the population of transit amplifying cells in more complex models. Models with and without feedback on the process of de-differentiation were considered. The same results were observed among the different models that were explored here. In contrast, the model by Shirayeh et al (Shirayeh et al., 2016) does not take into account feedback regulation, but assumed a constant population Moran process. Moreover, the models differ in their assumptions about the nature of stem cell divisions. Shirayeh et al assume both symmetric and asymmetric stem cell divisions, where symmetric stem cell division results in the generation of two progeny stem cells, and asymmetric division results in the generation of one progeny stem cell and one differentiated cell. Further work will have to determine which of the differences between the two modeling approaches gives rise to the different model predictions, and how this can be interpreted biologically.

Apart from the mutant invasion dynamics, our models suggest that another cancer-protecting mechanism can arise from the fact that in the presence of de-differentiation, two separate processes contribute to tissue growth and maintenance: stem cell self-renewal, and stem cell generation through de-differentiation. If each of these processes individually is not sufficient to drive uncontrolled growth in the absence of feedback, then feedback on both processes must be lost for disease to be initiated. Evolution of uncontrolled growth is thus less likely in the presence of both self-renewal and dedifferentiation compared to a situation where tissue is maintained only through stem cell self-renewal. This corresponds to a scenario where tissue maintenance is the result of division of labor between self-renewal and de-differentiation. If, however, the two processes are redundant, and tissue can be maintained by either self-renewal alone or by de-differentiation alone, then loss of homeostasis is more likely in the presence compared to the absence of de-differentiation. Loss of homeostasis due to the occurrence of cellular de-differentiation at a sufficient rate has also been reported mathematically by (Jilkine and Gutenkunst, 2014) in a different setting and modeling framework. In such a scenario, de-differentiation adds a second, independent, pathway towards uncontrolled growth, and a mutation resulting in feedback loss on either mechanism alone can drive uncontrolled cell growth, even if feedback on the second mechanism remains intact. Therefore, it will be important to quantify the rates of self-renewal and dedifferentiation in specific systems to determine whether they collaborate to ensure tissue maintenance, or whether they are redundant mechanisms.

## References

Anderson, A. R., Quaranta, V., 2008. Integrative mathematical oncology. Nat Rev Cancer 8, 227–34, doi:10.1038/nrc2329.

Cabrera, M. C., Hollingsworth, R. E., Hurt, E. M., 2015. Cancer stem cell plasticity and tumor hierarchy. World J Stem Cells 7, 27–36, doi:10.4252/wjsc.v7.i1.27.

Chaffer, C. L., Marjanovic, N. D., Lee, T., Bell, G., Kleer, C. G., Reinhardt, F., D'Alessio, A. C., Young, R. A., Weinberg, R. A., 2013. Poised chromatin at the ZEB1 promoter enables breast cancer cell plasticity and enhances tumorigenicity. Cell 154, 61–74, doi:10.1016/j.cell.2013.06.005.

Chaffer, C. L., Brueckmann, I., Scheel, C., Kaestli, A. J., Wiggins, P. A., Rodrigues, L. O., Brooks, M., Reinhardt, F., Su, Y., Polyak, K., Arendt, L. M., Kuperwasser, C., Bierie, B., Weinberg, R. A., 2011. Normal and neoplastic nonstem cells can spontaneously convert to a stem-like state. Proc Natl Acad Sci U S A 108, 7950–5, doi:10.1073/pnas.1102454108.

Desai, T. J., Brownfield, D. G., Krasnow, M. A., 2014. Alveolar progenitor and stem cells in lung development, renewal and cancer. Nature 507, 190–4, doi:10.1038/nature12930.

Dorantes-Acosta, E., Pelayo, R., 2012. Lineage switching in acute leukemias: a consequence of stem cell plasticity? Bone Marrow Res 2012, 406796, doi:10.1155/2012/406796.

Enderling, H., Hahnfeldt, P., 2011. Cancer stem cells in solid tumors: is 'evading apoptosis' a hallmark of cancer? Prog Biophys Mol Biol 106, 391–9, doi:10.1016/j.pbiomolbio.2011.03.007.

Enderling, H., Hlatky, L., Hahnfeldt, P., 2013. Cancer Stem Cells: A Minor Cancer Subpopulation that Redefines Global Cancer Features. Front Oncol 3, 76, doi:10.3389/fonc.2013.00076.

Enderling, H., Chaplain, M. A., Anderson, A. R., Vaidya, J. S., 2007. A mathematical model of breast cancer development, local treatment and recurrence. J Theor Biol 246, 245–59, doi:10.1016/j.jtbi.2006.12.010.

Ewens, W. J., 2004. Mathematical Population Genetics 1: Theoretical Introduction. Springer, New York.

Gillespie, D. T., 1976. General Method for Numerically Simulating Stochastic Time Evolution of Coupled Chemical-Reactions. Journal of Computational Physics 22, 403–434, doi:Doi 10.1016/0021-9991(76)90041-3.

Glauche, I., Cross, M., Loeffler, M., Roeder, I., 2007. Lineage specification of hematopoietic stem cells: mathematical modeling and biological implications. Stem Cells 25, 1791–9, doi:10.1634/stemcells.2007-0025.

Gupta, P. B., Fillmore, C. M., Jiang, G., Shapira, S. D., Tao, K., Kuperwasser, C., Lander, E. S., 2011. Stochastic state transitions give rise to phenotypic equilibrium in populations of cancer cells. Cell 146, 633–44, doi:10.1016/j.cell.2011.07.026.

Hartl, D. L., Clark, A. G., 1997. Principles of Population Genetics. Sinauer Associates, Sunderland, USA.

Huels, D. J., Sansom, O. J., 2015. Stem vs non-stem cell origin of colorectal cancer. Br J Cancer 113, 1–5, doi:10.1038/bjc.2015.214.

Jilkine, A., Gutenkunst, R. N., 2014. Effect of dedifferentiation on time to mutation acquisition in stem cell-driven cancers. PLoS Comput Biol 10, e1003481, doi:10.1371/journal.pcbi.1003481.

Jordan, C. T., Guzman, M. L., Noble, M., 2006. Cancer stem cells. N Engl J Med 355, 1253–61.

Kaveh, K., Kohandel, M., Sivaloganathan, S., 2016. Replicator dynamics of cancer stem cell: Selection in the presence of differentiation and plasticity. Math Biosci 272, 64–75, doi:10.1016/j.mbs.2015.11.012.

Komarova, N. L., van den Driessche, P., 2017. Stability of Control Networks in Autonomous Homeostatic Regulation of Stem Cell Lineages. Bull Math Biol, doi:10.1007/s11538-017-0283-4.

Konstorum, A., Hillen, T., Lowengrub, J., 2016. Feedback Regulation in a Cancer Stem Cell Model can Cause an Allee Effect. Bull Math Biol 78, 754–85, doi:10.1007/s11538-016-0161-5.

Kreso, A., Dick, J. E., 2014. Evolution of the cancer stem cell model. Cell Stem Cell 14, 275–91, doi:10.1016/j.stem.2014.02.006.

Kurtova, A. V., Xiao, J., Mo, Q., Pazhanisamy, S., Krasnow, R., Lerner, S. P., Chen, F., Roh, T. T., Lay, E., Ho, P. L., Chan, K. S., 2015. Blocking PGE2-induced tumour repopulation abrogates bladder cancer chemoresistance. Nature 517, 209–13, doi:10.1038/nature14034.

Lander, A. D., Gokoffski, K. K., Wan, F. Y., Nie, Q., Calof, A. L., 2009. Cell lineages and the logic of proliferative control. PLoS Biol 7, e15, doi:10.1371/journal.pbio.100001508-PLBI-RA-3368 [pii].

Leder, K., Holland, E. C., Michor, F., 2010. The therapeutic implications of plasticity of the cancer stem cell phenotype. PLoS One 5, e14366, doi:10.1371/journal.pone.0014366.

Li, Y., Laterra, J., 2012. Cancer stem cells: distinct entities or dynamically regulated phenotypes? Cancer Res 72, 576–80, doi:10.1158/0008-5472.CAN-11-3070.

Lo, W. C., Chou, C. S., Gokoffski, K. K., Wan, F. Y., Lander, A. D., Calof, A. L., Nie, Q., 2009. Feedback regulation in multistage cell lineages. Math Biosci Eng 6, 59–82.

Mani, S. A., Guo, W., Liao, M. J., Eaton, E. N., Ayyanan, A., Zhou, A. Y., Brooks, M., Reinhard, F., Zhang, C. C., Shipitsin, M., Campbell, L. L., Polyak, K., Brisken, C., Yang, J., Weinberg, R. A., 2008. The epithelial-mesenchymal transition generates cells with properties of stem cells. Cell 133, 704–15, doi:10.1016/j.cell.2008.03.027.

Marciniak-Czochra, A., Stiehl, T., Ho, A. D., Jager, W., Wagner, W., 2009. Modeling of asymmetric cell division in hematopoietic stem cells--regulation of self-renewal is essential for efficient repopulation. Stem Cells Dev 18, 377–85, doi:10.1089/scd.2008.0143.

Marjanovic, N. D., Weinberg, R. A., Chaffer, C. L., 2013. Cell plasticity and heterogeneity in cancer. Clin Chem 59, 168–79, doi:10.1373/clinchem.2012.184655.

Michor, F., 2008. Mathematical models of cancer stem cells. J Clin Oncol 26, 2854–61, doi:10.1200/jC0.2007.15.2421.

Nei, M., 1975. Molecular Population Genetics and Evolution. North-Holland Publishing Company, Amsterdam, Holland.

Nicolas, P., Kim, K. M., Shibata, D., Tavare, S., 2007. The stem cell population of the human colon crypt: analysis via methylation patterns. PLoS Comput Biol 3, e28, doi:10.1371/journal.pcbi.0030028.

Philpott, A., Winton, D. J., 2014. Lineage selection and plasticity in the intestinal crypt. Curr Opin Cell Biol 31, 39–45, doi:10.1016/j.ceb.2014.07.002.

Reya, T., Morrison, S. J., Clarke, M. F., Weissman, I. L., 2001. Stem cells, cancer, and cancer stem cells. Nature 414, 105–11.

Rodriguez-Brenes, I. A., Komarova, N. L., Wodarz, D., 2011. Evolutionary dynamics of feedback escape and the development of stem-cell-driven cancers. Proc Natl Acad Sci U S A 108, 18983–8, doi:10.1073/pnas.11076211081107621108 [pii].

Rodriguez-Brenes, I. A., Wodarz, D., Komarova, N. L., 2013a. Stem cell control, oscillations, and tissue regeneration in spatial and non-spatial models. Front Oncol 3, 82, doi:10.3389/fonc.2013.00082.

Rodriguez-Brenes, I. A., Komarova, N. L., Wodarz, D., 2013b. Tumor growth dynamics: insights into evolutionary processes. Trends Ecol Evol 28, 597–604, doi:10.1016/j.tree.2013.05.020.

Rodriguez-Brenes, I. A., Wodarz, D., Komarova, N. L., 2015. Characterizing inhibited tumor growth in stem-cell-driven non-spatial cancers. Math Biosci 270, 135–41, doi:10.1016/j.mbs.2015.08.009.

Rodriguez-Brenes, I. A., Kurtova, A. V., Lin, C., Lee, Y. C., Xiao, J., Mims, M., Chan, K. S., Wodarz, D., 2017. Cellular Hierarchy as a Determinant of Tumor Sensitivity to Chemotherapy. Cancer Res 77, 2231–2241, doi:10.1158/0008-5472.CAN-16-2434.

Roeder, I., Loeffler, M., 2002. A novel dynamic model of hematopoietic stem cell organization based on the concept of within-tissue plasticity. Exp Hematol 30, 853–61.

Roederer, I., Horn, M., Glauche, I., Hochhaus, A., Mueller, M. C., Loeffler, M., 2006. Leukemia stem cells - hit or miss. Nature Medicine Fill in.

Scheel, C., Weinberg, R. A., 2011. Phenotypic plasticity and epithelial-mesenchymal transitions in cancer and normal stem cells? Int J Cancer 129, 2310–4, doi:10.1002/ijc.26311.

Shirayeh, A. M., Kaveh, K., Kohandel, M., Sivaloganathan, S., 2016. Phenotypic heterogeneity in modeling cancer evolution. arXiv arXiv:1610.08163.

Simons, B. D., Clevers, H., 2011. Strategies for homeostatic stem cell self-renewal in adult tissues. Cell 145, 851–62, doi:10.1016/j.cell.2011.05.033.

Snippert, H. J., van der Flier, L. G., Sato, T., van Es, J. H., van den Born, M., Kroon-Veenboer, C., Barker, N., Klein, A. M., van Rheenen, J., Simons, B. D., Clevers, H., 2010. Intestinal crypt homeostasis results from neutral competition between symmetrically dividing Lgr5 stem cells. Cell 143, 134–44, doi:10.1016/j.cell.2010.09.016.

Stange, D. E., Koo, B. K., Huch, M., Sibbel, G., Basak, O., Lyubimova, A., Kujala, P., Bartfeld, S., Koster, J., Geahlen, J. H., Peters, P. J., van Es, J. H., van de Wetering, M., Mills, J. C., Clevers, H., 2013. Differentiated Troy+ chief cells act as reserve stem cells to generate all lineages of the stomach epithelium. Cell 155, 357–68, doi:10.1016/j.cell.2013.09.008.

Stiehl, T., Marciniak-Czochra, A., 2011. Characterization of stem cells using mathematical models of multistage cell lineages. Mathematical and Computer Modelling https://doi.org/10.1016/j.mcm.2010.03.057.

Sun, Z., Komarova, N. L., 2015. Stochastic control of proliferation and differentiation in stem cell dynamics. J Math Biol 71, 883–901, doi:10.1007/s00285-014-0835-2.

Tata, P. R., Mou, H., Pardo-Saganta, A., Zhao, R., Prabhu, M., Law, B. M., Vinarsky, V., Cho, J. L., Breton, S., Sahay, A., Medoff, B. D., Rajagopal, J., 2013. Dedifferentiation of committed epithelial cells into stem cells in vivo. Nature 503, 218–23, doi:10.1038/nature12777.

Tonekaboni, S. A., Dhawan, A., Kohandel, M., 2017. Mathematical modelling of plasticity and phenotype switching in cancer cell populations. Math Biosci 283, 30–37, doi:10.1016/j.mbs.2016.11.008.

Visvader, J. E., Lindeman, G. J., 2012. Cancer stem cells: current status and evolving complexities. Cell Stem Cell 10, 717–28, doi:10.1016/j.stem.2012.05.007.

Watt, F. M., Hogan, B. L., 2000. Out of Eden: stem cells and their niches. Science 287, 1427–30.

Weissman, I. L., 2000. Stem cells: units of development, units of regeneration, and units in evolution. Cell 100, 157–68.

Werner, B., Dingli, D., Lenaerts, T., Pacheco, J. M., Traulsen, A., 2011. Dynamics of mutant cells in hierarchical organized tissues. PLoS Comput Biol 7, e1002290, doi:10.1371/journal.pcbi.1002290.

Werner, B., Scott, J. G., Sottoriva, A., Anderson, A. R., Traulsen, A., Altrock, P. M., 2016. The Cancer Stem Cell Fraction in Hierarchically Organized Tumors Can Be Estimated Using Mathematical Modeling and Patient-Specific Treatment Trajectories. Cancer Res 76, 1705–13, doi:10.1158/0008-5472.CAN-15-2069.

Yang, J., Plikus, M. V., Komarova, N. L., 2015. The Role of Symmetric Stem Cell Divisions in Tissue Homeostasis. PLoS Comput Biol 11, e1004629, doi:10.1371/journal.pcbi.1004629.

Yanger, K., Stanger, B. Z., 2014. Liver cell reprogramming: parallels with iPSC biology. Cell Cycle 13, 1211–2, doi:10.4161/cc.28381.

Yanger, K., Zong, Y., Maggs, L. R., Shapira, S. N., Maddipati, R., Aiello, N. M., Thung, S. N., Wells, R. G., Greenbaum, L. E., Stanger, B. Z., 2013. Robust cellular reprogramming occurs spontaneously during liver regeneration. Genes Dev 27, 719–24, doi:10.1101/gad.207803.112.

